# Pharmacogenetic allele variant frequencies: An analysis of the VA’s Million Veteran Program (MVP) as a representation of the diversity in US population

**DOI:** 10.1101/2022.08.26.505483

**Authors:** K Markianos, F Dong, B Gorman, Y Shi, D Dochterman, U Saxena, P Devineni, J Moser, S Muralidhar, R Ramoni, P Tsao, Million Veteran Program, S Pyarajan, R Przygodzki

## Abstract

We present allele frequencies of pharmacogenomics relevant variants across multiple ancestry in a sample representative of the US population. We analyzed 658,582 individuals with genotype data and extracted pharmacogenomics relevant single nucleotide variant (SNV) alleles, human leukocyte antigens (HLA) 4-digit alleles and an important copy number variant (CNV), the full deletion/duplication of CYP2D6. We compiled distinct allele frequency tables for European, African American, Hispanic, and Asian ancestry individuals. In addition, we compiled allele frequencies based on local ancestry reconstruction in the African-American (2-way deconvolution) and Hispanic (3-way deconvolution) cohorts.

## Introduction

Genetic polymorphisms of metabolic pathways and cytochrome P450 (CYP) genes are associated with altering pharmacokinetics and pharmacodynamics of the absorption, distribution, metabolism and excretion (ADME) of drug and toxic compounds (xenobiotics). Gaining a better understanding of the interindividual variations of this genetic makeup is necessary to understand the metabolic rate of efficiency a xenobiotic is metabolized. In general, heritable selective pressure is a major determinant of variant frequency among the different ethnic populations, typically presenting with two or more variants identified in most metabolic pathway genes. Common star allelic variants (referred to herein as “variant”) prescribe a “normal” metabolic cycle while others convey a heightened or depressed metabolic cycle. Much of this is well catalogued in a variety of collections, including PharmGKB **(https://www.pharmgkb.org/)**, with clinically actionable variant-vs-drug combinations presented in the Clinical Pharmacogenetics Implementation Consortium **(CPIC https://cpicpgx.org/)** and the Dutch Pharmacogenetics Working Group **(DPWG http://upgx.eu/guidelines)**. While these variants are catalogued in the multitude of databases, it is also important to recognize that many of the variants identified heavily rely upon data derived from unique ethnic populations. Ethnic population data are typically derived from a limited collection of self-identified subjects and the unique variants associated within that ethnicity. Moreover, certain variants designated as normal are unique to a select ethnicity and not represented among others, such as is known for CYP2D6 and codeine, or CYP3A5 and Tacrolimus (1). Lastly, while certain populations are considered relatively homogenous over several generations as dictum of culture, not all data is reflective of this consideration which further contributes to the diversity of drug responses.

The distribution of inherited xenobiotic-metabolizing alleles differs considerably between populations (2, 3) and appears to be rigid in frequency among ethnically stable populations. While there are several large-scale data sets that can provide variant frequencies of pharmacogenomic genes for researchers and clinicians to use, most of these data are not representative of the “melting pot” of the genetic ancestries present within the United States. (US). While one could rely on self-reported ethnicity to improve variant frequency found among unique populations, such data is imperfect (4). Further complicating possible variant predictions is that nearly everyone has at least one pharmacogenomic variant allele with as many as 3% carrying 5 allelic variants (5). These findings limit the overarching use of ethnically related variant frequencies in diverse populations such as is present in the US. This is because the data available is limited to a specific self-reported ethnicity and/or does not consider other variants that could be present among other ethnicities. This is a particularly important consideration for research of personalized drug therapy and potentially changes the healthcare guidelines provided by groups such as CPIC and DPWG that select important alleles for clinical genotyping based in part on population prevalence.

To address the issues with the use of pharmacogenomic variants and to further explore possibly pharmacogenetically-associated variant markers we used the Million Veteran Program (MVP) (6) with >800,000 participants to generate a coherent representation of allelic frequencies present within a US population. The MVP cohort is mostly male but is very diverse and represents the US population ancestry in general. The genotype data was imputed using the African Genome Resources (AGR) and 1000 Genomes imputation reference panels.

## Results

Our analysis is based on the Release 4 of MVP data with 658,582 individuals genotyped with the MVP-1 Axiom array(7). Participants were assigned ancestry based on the HARE algorithm (8). The MVP cohort is diverse with ∼30% of the cohort assigned as non-European (EUR 467k, AFR 125k, HIS 52k, ASN 8k). A small fraction of the cohort was highly admixed and not assigned to any of the four major ancestries and is not included in this analysis (<2%).

Our aim is to provide ancestry specific variant frequency catalog for a significant fraction of pharmacogenomics relevant variants in a large cohort representative of the US population. We examined Single Nucleotide Variants (SNV), an important pharmacogenetics relevant Copy Number Variant (CNV) as well as Human Leukocyte Antigen (HLA) 4-digit alleles. We defined our pharmacogenomics gene set by combining information from two publicly available data bases, PharmGKB and PharmVar (Methods). Overall, we were able to determine SNV frequencies for 273/1339 targeted SNVs, in 148/152 targeted genes. Details on variant selection can be found in Methods. As expected, SNV allele frequencies vary substantially among HARE groups (Figure 1).

**Figure 1.**
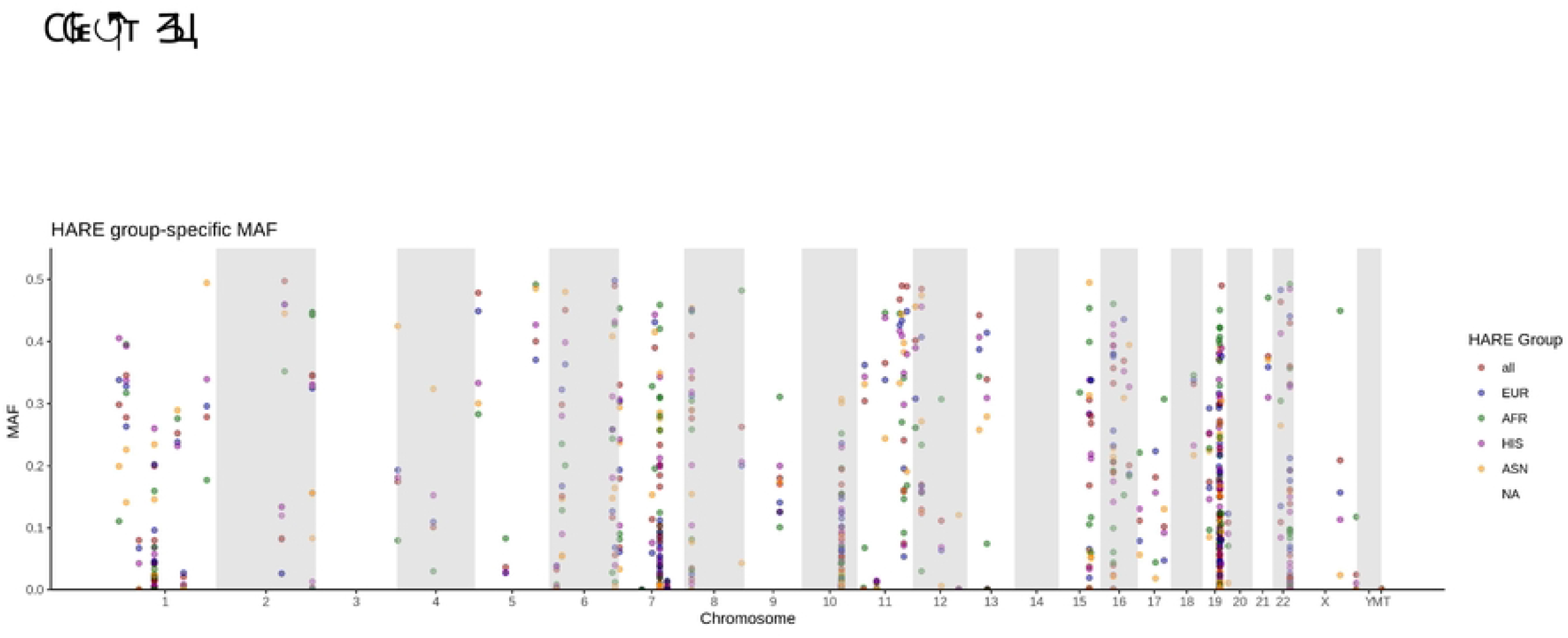
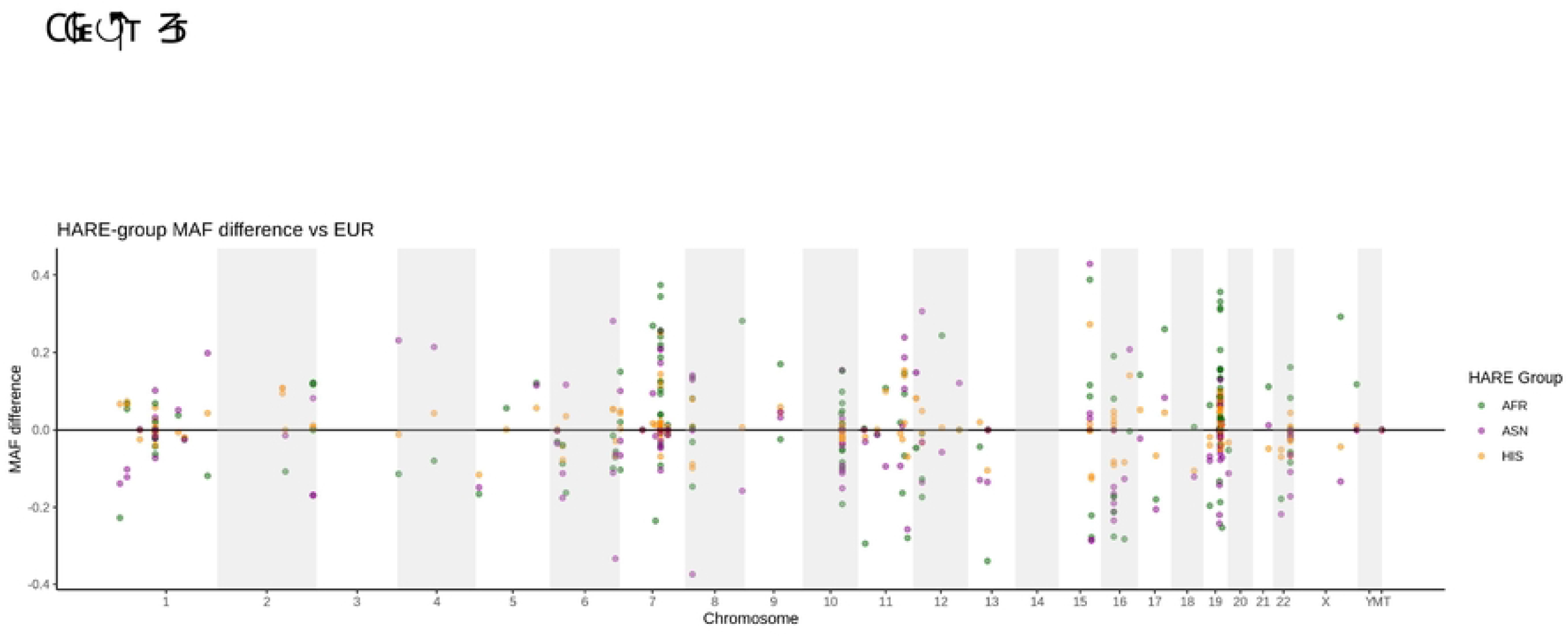
(a) Allele frequency distributions in different ethnic groups of 282 variants in 153 genes (b) differences in allele frequencies relative to EUR samples.

In addition to HARE allele frequencies, we used Local Ancestry Inference (LAI) to identify ancestral origin of individual chromosomal segments and compute allele frequencies based on the local ancestry. We “painted” the African American samples (125 k individuals) using two-way deconvolution, extracting allele frequencies for the AFR and EUR tracks. For the Hispanic individuals (52 k) we used three-way deconvolution to compute allele frequencies for EUR, AFR and Native American (AMR) tracks. Details on the LAI will be presented elsewhere. In Figure 2 we present allele frequencies derived from HARE groups, LAI as well as two publicly available databases, 1 k genome and gnomAD (Methods). The most striking differences are observed for Hispanics, a group that is extremely heterogeneous and not well defined in the genetics literature.

**Figure 2.**
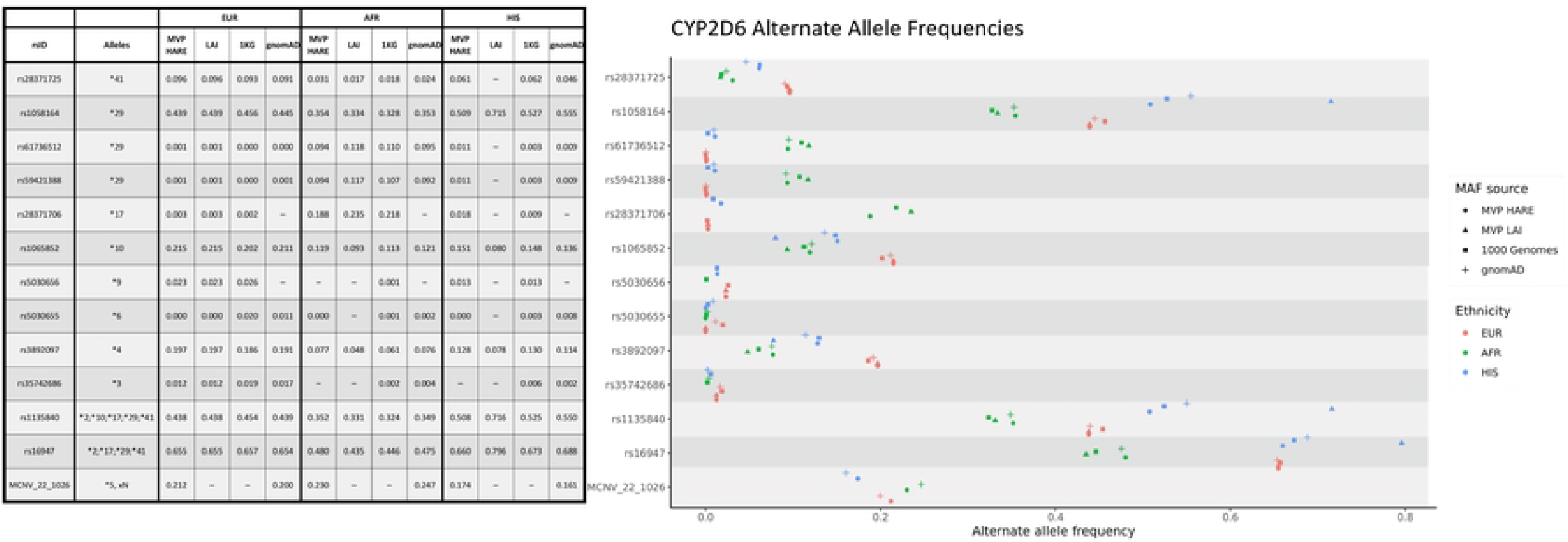
CYP2D6 allele frequencies for three MVP HARE groups, MVP Local Ancestry Inference (LAI) and two publicly available data sets: 1000 Genome Project and gnomAD. Only consensus Tier 1 alleles (9) are shown.

Allele frequencies for the three major MVP HARE groups (EUR, AFR, HIS) are in good agreement with gnomAD derived estimates. However, comparison of gnomAD HIS frequencies with the LAI AMR track of the MVP HIS population reveals significant differences (Supplementary Figure 1). Here we note that the LAI AMR track provides much better allele frequency ascertainment than the 60 AMR genomes that were used to anchor the local ancestry deconvolution. In the MVP HARE HIS population (52k individuals) the AMR track contributes ∼30% of the genome resulting in an effective population size of ∼15k individuals. Furthermore, while the 60 AMR genomes we used to anchor ancestry deconvolution provide sufficient multi-locus information to resolve local ancestry, the AMR track is a much better sampling of the AMR genome as it exists today in the US population.

Sirolimus is a widely used immunosuppressant and the variant controlling its metabolism, rs2242480 of CYP3A4 (10), varies among populations. We observe widely different allele frequencies in the three major groups (EUR, AFR, HIS) for gnomAD (0.09, 0.74, 0.37) and MVP (0.09, 0.73, 0.35). However, the local ancestry derived AMR allele frequency (0.67) is almost twice as high as the HIS allele frequency. The same observation applies to Tacrolimus, another significant immunosuppressant. The controlling variant, rs776746 of CYP3A5, shows large variation in major MVP groups (0.07, 0.70, 0.21) and there is a significant difference between HIS and local ancestry derived AMR allele frequency (0.31). Thus, recent demographic history of individuals, and the fraction of inheritance derived from different major population groups, has a large impact on the allele frequency distribution of pharmacogenomics relevant variants.

CYP2D6 is an important component of cytochrome P450 and is involved in the metabolism of many commonly prescribed medications, including antidepressants, antipsychotics, b-blockers, opioids, antiemetics, atomoxetine, and tamoxifen (9, 11). In addition to SNV frequencies for the most significant variants (12), Figure 2 presents allele frequencies for an important copy number variant, the whole gene deletion designated as CYP2D6*5 in the pharmacogenomics literature (Figures 2 and 3). We called the CYP2D6*5 CNV using UMAP, a machine learning algorithm (13). Assignments are clearly separated for copy gain and copy loss. Furthermore, we can clearly separate single and double copy loss (Figure 3). The UMAP approach offers a clear advantage over PCA based classification (Supplementary Figure 2).

**Figure 3.**
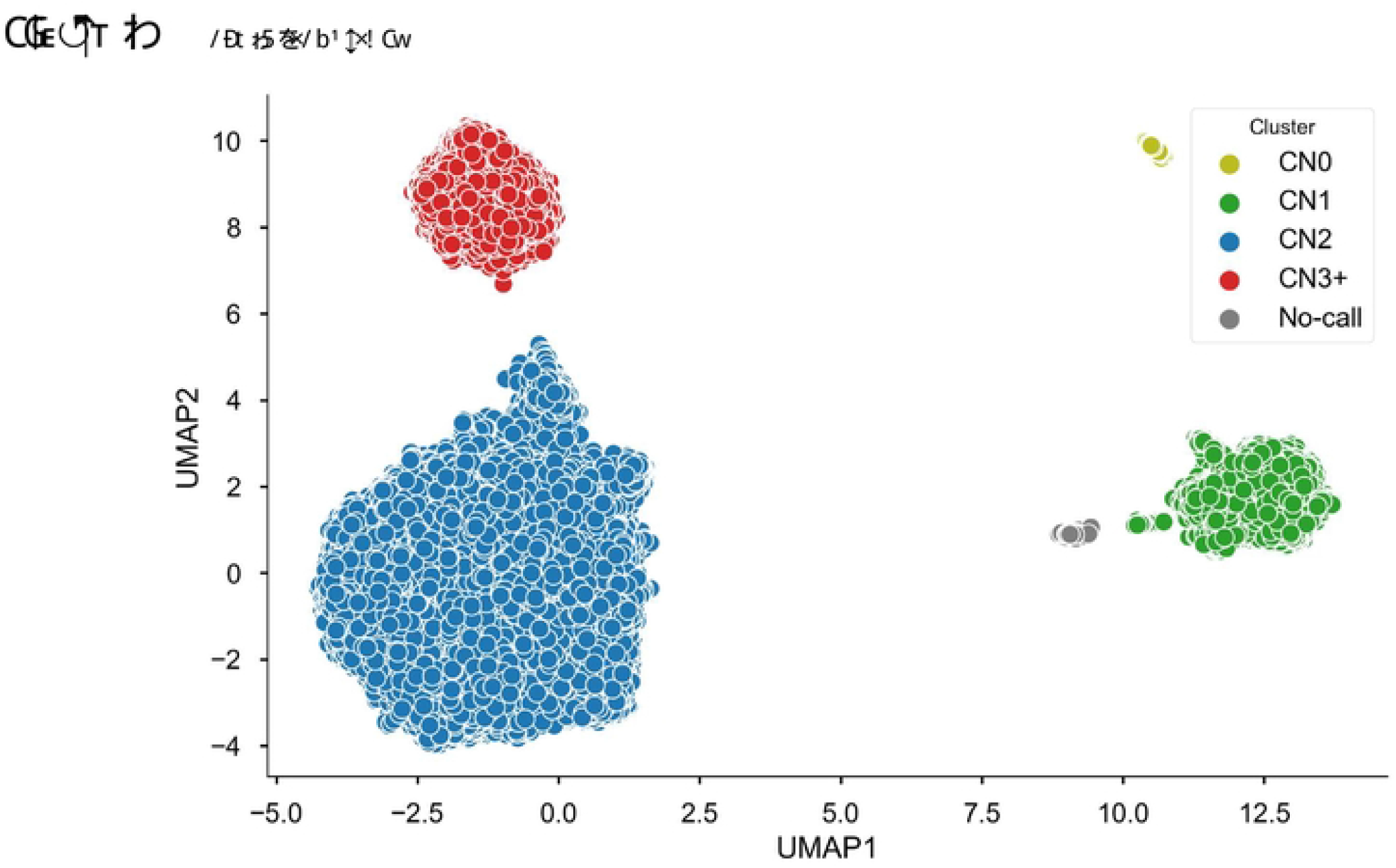
Copy number variation in CYP2D6. Results are shown just for the HARE AFR cohort; clusters were derived using UMAP (13).

We note that UMAP does not represent a general approach to copy number variation detection. Hyper-parameters for the model are tuned for the specific, relatively common CNVs. Furthermore, we achieve optimal performance only when we tune the model separately for individual HARE ancestries.

**Supplementary Figure 2.**
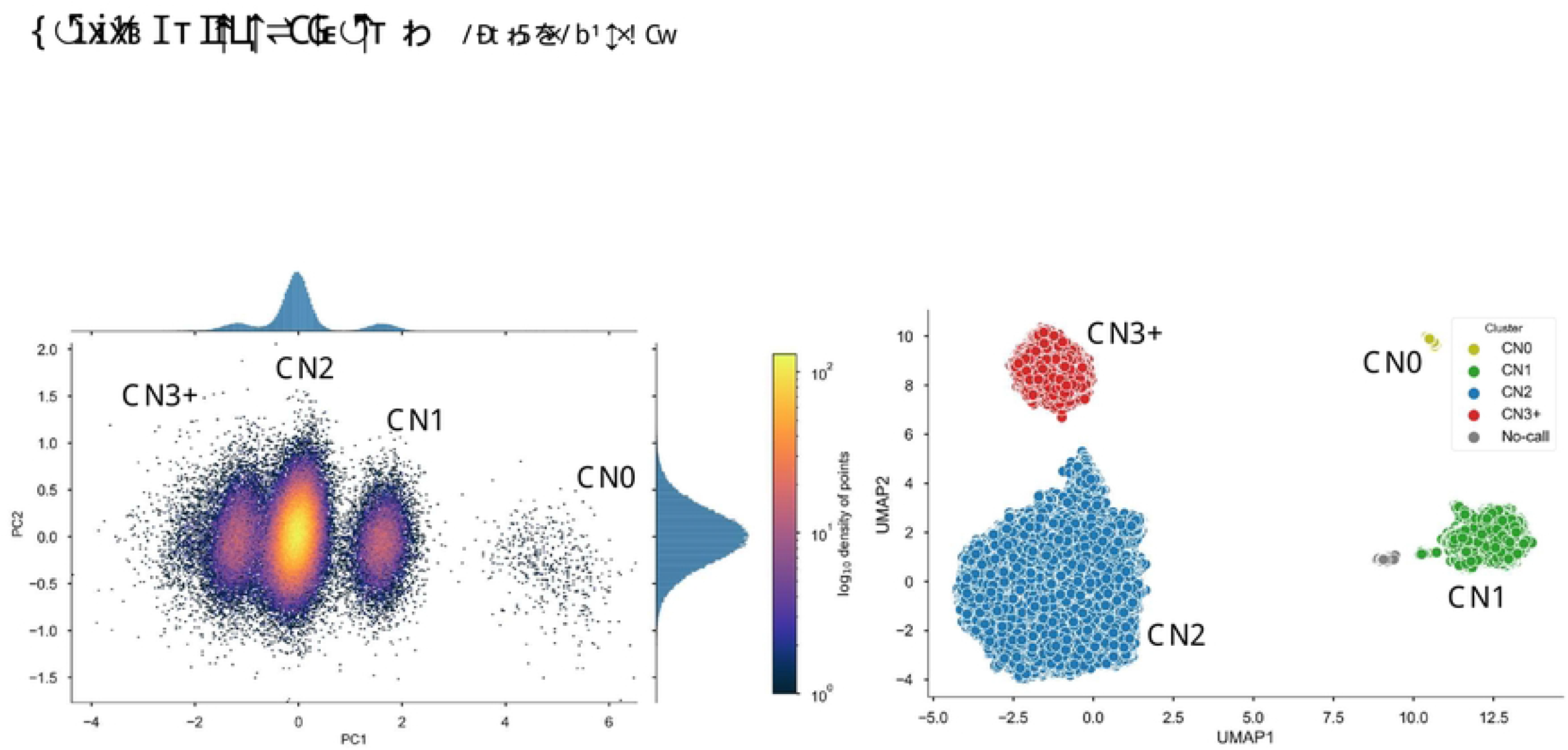
Copy number variation in CYP2D6 using two computational approaches. Results are shown just for the HARE AFR cohort; clusters were derived using (a) Principal Components Analysis (PCA) and (b) UMAP(13). UMAP significantly reduces assignment ambiguity.

Table 1a presents CYP2D6*5 allele frequencies for the three major HARE groups. The major survey of CYP2D6*5 (14) finds slightly different allele frequencies, e.g., 89% in Beoris et al vs 78% in MVP for copy number 2 EUR. Our findings are closer to the frequencies reported by gnomAD (80%, MCNV_22_1026 | gnomAD SVs v2.1 | gnomAD (broadinstitute.org). The differences might be due to different assays: single site PCR vs SNP genotyping (MVP) or sequencing (gnomAD), or differences in ascertainment of ethnic background. We are not able to run the UMAP algorithm on phased chromosomes, but we can use ancestry deconvolution and test CNV status in individuals ancestry-homozygous at CYP2D6, e.g. AMR/AMR individuals in the HARE HIS group. Results are shown in Table 1b. The ancestral AMR genome clearly harbors a lot fewer non-diploid samples while the EUR tracks are in close agreement with observations in the major HARE groups.

**Table 1.** Allele frequencies for allele CYP2D6*5 (whole gene deletion) in different HARE groups. In addition, we show allele frequencies in the three components of the HARE HIS group (EUR, AFR, AMR) calculated using individuals ancestry-homozygous at CYP2D6.

| Copies | HARE group |  |  | HARE HIS, ancestry homozygous segments |  |  |
| --- | --- | --- | --- | --- | --- | --- |
|  | EUR | AFR | HIS | EUR | AFR | AMR |
| 0 | 0.14 | 0.37 | 0.14 | 0.16 | 0.31 | 0.03 |
| 1 | 6.25 | 10.70 | 5.71 | 5.93 | 9.87 | 4.47 |
| 2 | 78.40 | 76.90 | 82.50 | 77.01 | 79.78 | 93.05 |
| 3+ | 14.80 | 11.96 | 11.58 | 16.89 | 10.03 | 2.45 |

In addition to SNV and specific CNVs we derived HLA alleles from SNP genotypes using the HIBAG algorithm (15). Although HLA status does not modify pharmacokinetics there are well established adverse drug reactions in the presence of specific HLA alleles. For example, abacavir, a common anti-retroviral, causes abacavir hypersensitivity syndrome in the presence of HLA-*5701; Allopurinol is typically a safe drug for the treatment of gout but in the presence of HLA-B*58:01 is associated with an increased risk for allopurinol induced SCAR and SJS/TEN (16). HLA 4-digit Class I and Class II allele distribution for four HARE groups is shown in Figure 4. As expected, allele frequencies are highly variable in the four groups, including the three alleles most relevant for pharmacogenomics: HLA-A*3101, HLA-B*5701 and HLA-B*58:01 (Table 2). Details of HLA allele imputation will be presented elsewhere. Here, we note that HLA imputation precision was >90% for HARE EUR, AFR and HIS groups. However, we currently observe lower precision for the ASN predictions due to lack of an appropriate training set, something we hope to address in the future.

**Figure 4.**
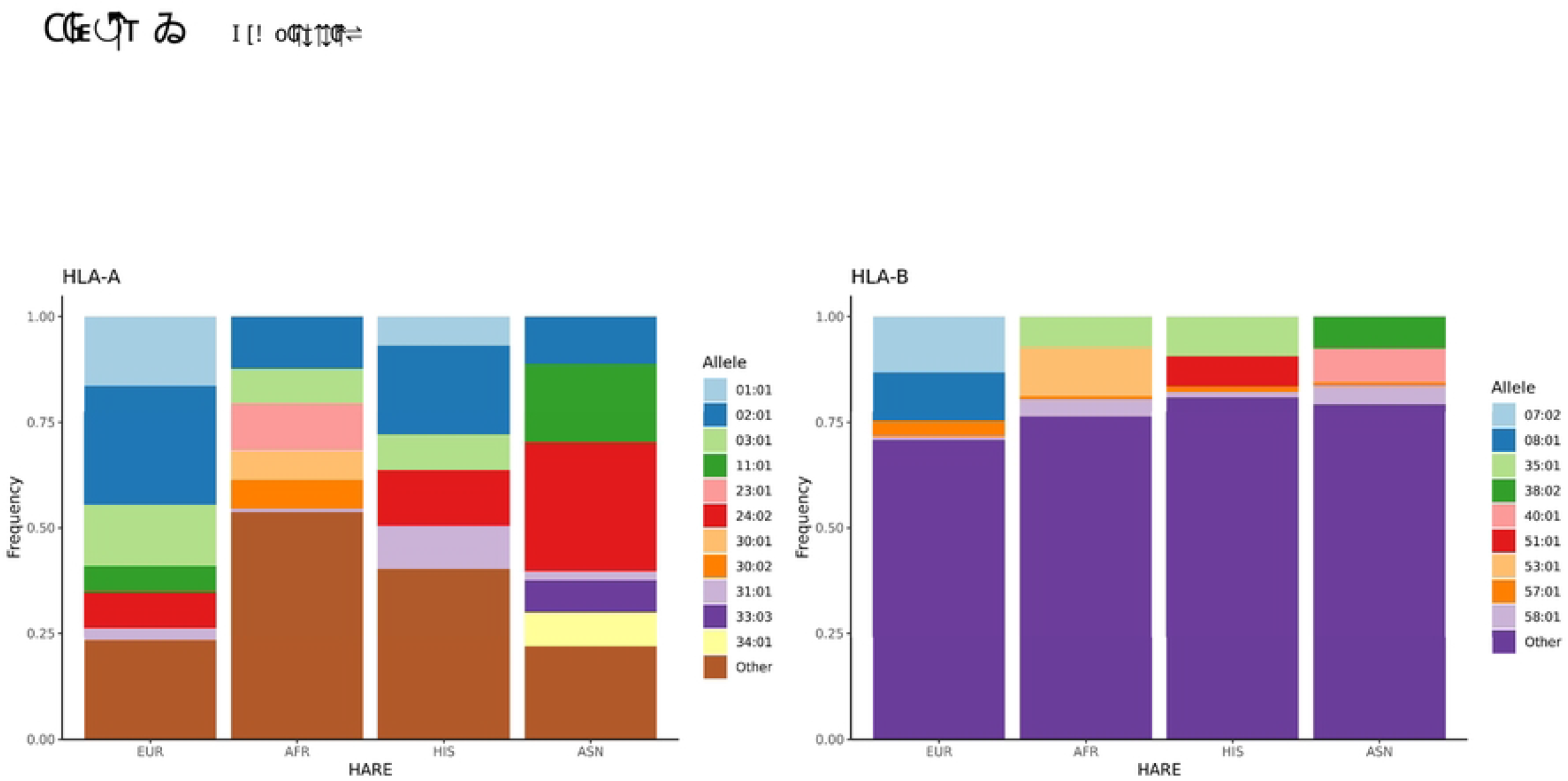
HLA allele distribution in 4 ethnic groups. Allele imputation was performed through HIBAG using the Axiom UK Biobank model and Axiom MVP genotypes.

**Table 2.** Allele frequencies for HLA alleles with known large effects in pharmacogenomics.

| Allele | EUR | AFR | HIS | ASN |
| --- | --- | --- | --- | --- |
| HLA-A*31:01 | 2.75 | 0.82 | 5.08 | 2.21 |
| HLA-B*57:01 | 3.73 | 0.79 | 1.41 | 0.75 |
| HLA-B*58:01 | 0.68 | 4.07 | 1.23 | 4.45 |

## Discussion

We present a survey of pharmacogenetics relevant variants in the MVP, a sample representative of the US population. Using the MVP-1 Axiom array we can resolve a large fraction of known pharmacogenomics alleles, either as direct or as imputed genotypes. In addition, we use the genotypes to derive population allele frequencies for an important common CNV, whole gene deletion/duplication of CYP2D6, as well as population distribution of HLA alleles, including HLA alleles important for drug delivery decisions.

As expected, there is substantial variation in allele frequencies between ancestry groups for a subset of the examined variants. In addition, we show through local ancestry painting that in admixed populations such as African Americans and Hispanics we can see substantial differences in allele frequency dependent on ancestral origin of specific chromosomal sections or haplotypes. This observation further underscores the need for individual typing rather than reliance on self-reported ethnicity on drug delivery decisions in clinical practice.

There are limitations in our derived population allele frequencies. While SNVs are phased neither CNV nor HLA calls are phased genotypes. Furthermore, successful phasing in the overall genome does not guarantee successful phasing in complex genomic regions such as CYP2D6. For CYP2D6 in particular, we have been able to resolve whole gene deletions and duplications, but we are certain that there is additional small scale copy number variation that cannot be resolved by our UMAP machine learning approach. For example, small deletions and complex rearrangements involving the proximal CYP2D7 and CYP2D8 pseudogenes. It is likely that such complex variation has a minor contribution to the population distribution of CYP2D6 pharmacogenetics. However, resolution of population level frequency of such variants will require specialized assays such as long-range sequencing. Improving phasing and imputation will aid the eventual derivation of star alleles in these regions.

We think that this comprehensive allele frequency report in a population representative of the US genome diversity will become a useful reference for future guidelines of relative importance of alleles worth ascertaining in pharmacogenetics screens.

## Acknowledgements

This research is based on data from the Million Veteran Program, Office of Research and Development, Veterans Health Administration and was supported by award no. MVP000. This publication does not represent the views of the Department of Veterans Affairs, the US Food and Drug Administration, or the US Government.

## Consortia

VA Million Veteran Program

MVP Executive Committee

- Co-Chair: J. Michael Gaziano, M.D., M.P.H.
- Co-Chair: Rachel Ramoni, D.M.D., Sc.D.
- Kyong-Mi Chang, M.D.
- Grant Huang, Ph.D.
- Sumitra Muralidhar, Ph.D.
- Philip S. Tsao, Ph.D. MVP Program Office
- Sumitra Muralidhar, Ph.D.
- Jennifer Moser, Ph.D. MVP Recruitment/Enrollment
- Recruitment/Enrollment Director/Deputy Director, Boston
- Stacey B. Whitbourne, Ph.D.; Jessica V. Brewer, M.P.H.
- MVP Coordinating Centers
  - Clinical Epidemiology Research Center (CERC), West Haven – John Concato, M.D., M.P.H.
  - Cooperative Studies Program Clinical Research Pharmacy Coordinating Center, Albuquerque - Stuart Warren, J.D., Pharm D.; Dean P. Argyres, M.S.
  - Genomics Coordinating Center, Palo Alto – Philip S. Tsao, Ph.D.
  - Massachusetts Veterans Epidemiology Research Information Center (MAVERIC), Boston - J. Michael Gaziano, M.D., M.P.H.
  - MVP Information Center, Canandaigua – Brady Stephens, M.S.
- Core Biorepository, Boston – Mary T. Brophy M.D., M.P.H.; Donald E. Humphries, Ph.D.
- MVP Informatics, Boston – Nhan Do, M.D.; Shahpoor Shayan
- Center for Data and Computational Science (C-DACS) – Saiju Pyarajan, Ph.D.
- Data Operations/Analytics, Boston – Xuan-Mai T. Nguyen, Ph.D. MVP Science
- Genomics - Saiju Pyarajan Ph.D.; Philip S.Tsao, Ph.D.
- Phenomics - Kelly Cho, M.P.H, Ph.D.
- Data and Computational Sciences – Saiju Pyarajan, Ph.D.
- Statistical Genetics – Elizabeth Hauser, Ph.D.; Yan Sun, Ph.D.; Hongyu Zhao, Ph.D. MVP Local Site Investigators
- Atlanta VA Medical Center (Peter Wilson) - Bay Pines VA Healthcare System (Rachel McArdle)
- Birmingham VA Medical Center (Louis Dellitalia)
- Cincinnati VA Medical Center (John Harley)
- Clement J. Zablocki VA Medical Center (Jeffrey Whittle)
- Durham VA Medical Center (Jean Beckham)
- Edith Nourse Rogers Memorial Veterans Hospital (John Wells)
- Edward Hines, Jr. VA Medical Center (Salvador Gutierrez)
- Fayetteville VA Medical Center (Gretchen Gibson)
- VA Health Care Upstate New York (Laurence Kaminsky)
- New Mexico VA Health Care System (Gerardo Villareal)
- VA Boston Healthcare System (Scott Kinlay)
- VA Western New York Healthcare System (Junzhe Xu)
- Ralph H. Johnson VA Medical Center (Mark Hamner)
- Wm. Jennings Bryan Dorn VA Medical Center (Kathlyn Sue Haddock)
- VA North Texas Health Care System (Sujata Bhushan)
- Hampton VA Medical Center (Pran Iruvanti)
- Hunter Holmes McGuire VA Medical Center (Michael Godschalk)
- Iowa City VA Health Care System (Zuhair Ballas)
- Jack C. Montgomery VA Medical Center (Malcolm Buford)
- James A. Haley Veterans’ Hospital (Stephen Mastorides)
- Louisville VA Medical Center (Jon Klein)
- Louis Stokes Cleveland VA Medical Center (Frank Jacono)
- Manchester VA Medical Center (Nora Ratcliffe)
- Miami VA Health Care System (Hermes Florez)
- Michael E. DeBakey VA Medical Center (Alan Swann)
- Minneapolis VA Health Care System (Maureen Murdoch)
- N. FL/S. GA Veterans Health System (Peruvemba Sriram)
- Northport VA Medical Center (Shing Shing Yeh)
- Overton Brooks VA Medical Center (Ronald Washburn)
- Philadelphia VA Medical Center (Darshana Jhala)
- Phoenix VA Health Care System (Samuel Aguayo)
- Portland VA Medical Center (David Cohen)
- Providence VA Medical Center (Satish Sharma)
- Richard Roudebush VA Medical Center (John Callaghan)
- Salem VA Medical Center (Kris Ann Oursler)
- San Francisco VA Health Care System (Mary Whooley)
- South Texas Veterans Health Care System (Sunil Ahuja)
- Southeast Louisiana Veterans Health Care System (Amparo Gutierrez)
- Southern Arizona VA Health Care System (Ronald Schifman)
- Sioux Falls VA Health Care System (Jennifer Greco)
- St. Louis VA Health Care System (Michael Rauchman)
- Syracuse VA Medical Center (Richard Servatius)
- VA Eastern Kansas Health Care System (Mary Oehlert)
- VA Greater Los Angeles Health Care System (Agnes Wallbom)
- VA Loma Linda Healthcare System (Ronald Fernando)
- VA Long Beach Healthcare System (Timothy Morgan)
- VA Maine Healthcare System (Todd Stapley)
- VA New York Harbor Healthcare System (Scott Sherman)
- VA Pacific Islands Health Care System (Gwenevere Anderson)
- VA Palo Alto Health Care System (Philip Tsao)
- VA Pittsburgh Health Care System (Elif Sonel)
- VA Puget Sound Health Care System (Edward Boyko)
- VA Salt Lake City Health Care System (Laurence Meyer)
- VA San Diego Healthcare System (Samir Gupta)
- VA Southern Nevada Healthcare System (Joseph Fayad)
- VA Tennessee Valley Healthcare System (Adriana Hung)
- Washington DC VA Medical Center (Jack Lichy)
- W.G. (Bill) Hefner VA Medical Center (Robin Hurley)
- White River Junction VA Medical Center (Brooks Robey)
- William S. Middleton Memorial Veterans Hospital (Robert Striker)

## Author Contributions

PS and PR conceived and directed research, MK and DF performed data analysis, MK coordinated research efforts, GB contributed to data analysis and developed the UMAP CNV prediction model, SY DD SU and DP contributed to data analysis, MS RR and PT provided coordination and organizational support, MK DF GB PS and PR wrote and revised the manuscript.

## Methods

### Ethics statement

The Veterans Affairs (VA) central institutional review board (cIRB) and site-specific IRBs approved the Million Veteran Program study.

### MVP genotype data

The MVP Release 4 dataset includes 658,582 individuals and consists of a hard-called dataset of 667,955 variants prepared as described in Hunter-Zinck et al. 2020 (7), as well as an imputed dataset. QC’d genotypes were further prepared for phasing and imputation by removing markers with high missingness (>20%), monomorphic markers, and markers significantly out of Hardy-Weinberg equilibrium (p < 1e-6 adjusted for ancestry). Haplotypes were then statistically phased using SHAPEIT v4.1.3 and imputed into the African Genome Resources and 1000 Genomes imputation panels using Minimac4. Each individual in the cohort was assigned a HARE group (EUR, AFR, HIS, or ASN), a surrogate variable for ancestry and race/ethnicity (Fang et al. 2019). The MVP Release 4 cohort consists of 467,162 EUR, 124,756 AFR, 52,423 HIS, 8,364 ASN, and 5,877 unassigned individuals. All data are based in GRCh37.

### Identification of known pharmacogenetics variants

We curated a catalogue of known or high-confidence pharmacogenetics variants by rsID from the PharmGKB and PharmVar databases. From PharmGKB, we downloaded variant summary data

(https://api.pharmgkb.org/v1/download/file/data/variants.zip) and kept only variants with at least one Level 1 or 2 PharmGKB clinical annotation. From PharmVar, we downloaded the complete database (version 4.2.4) and kept all variants. In total, we identify 1,339 unique variants from 152 genes.

### Identification of pharmacogenetics variants in the MVP genotype dataset

#### Genotyped dataset

We selected the intersection of known pharmacogenetics variants with the catalog of SNPs in the MVP array. We identified pharmacogenetics variants by chromosome location and rsID (Hunter-Zinck et al., 2020).

#### Imputed dataset

Imputation was performed using MINIMAC. We kept only variants with imputation R^2^ > 0.9 within the ethnic group. We assigned rsIDs to imputed variants by intersecting variant genomic position with rsID genomic position in NCBI dbSNP (v154) using bedtools. We then identified pharmacogenetics variants by overlapping imputed variant rsIDs.

In total, we find 193 pharmacogenetics variants from 136 genes in the genotyped data set. Including the imputed variants, we expand the set to 282 variants in 153 genes.

### Allele frequency analyses

#### Calculation of minor allele frequencies

HARE group-specific minor allele frequencies (MAFs) were calculated for the MVP hard-called and imputed datasets using PLINK2.

#### Local Ancestry Inference (LAI) based allele frequencies

Briefly, we performed LAI using rfmix2. We used 3,942 reference samples for EUR, AFR and Native American (AMR) ancestry collected by the 1000 genome project and the Human Genome Diversity Project (HGDP). The reference VCF files were curated by the gnomAD team (https://gnomad.broadinstitute.org/downloads#v3-hgdp-1kg). We used local ancestry output to create separate, ancestry specific, VCF output files. Two files for the HARE AFR sample (EUR-AFR) and three files for the HARE Hispanic sample (EUR-AFR-NAT). The allele frequency extraction procedure was the same for LAI and gnomAD samples, described below.

#### GnomAD allele frequencies

Population-specific frequencies were extracted as follows. LAI and gnomAD (v2.1.1 Genomes only, not Exomes) frequencies were stored in the INFO fields of VCF files. AFR and HIS LAI frequencies were stored in separate files. gnomAD frequencies were stored in population-specific INFO fields (AF_nfe, AF_afr, AF_amr for non-Finnish Europeans, African/African Americans, and Latino/Admixed Americans respectively). Using bcftools 1.10, VCF files were first filtered to the relevant SNPs (bcftools view --include ‘ID=@<file of rsIDs> ’ <VCF file>), and frequencies were then extracted from the relevant INFO fields (e.g., bcftools query - ‘%ID\t%INFO/AF_nfe\n’).

#### 1000 genomes allele frequencies

1000 Genomes population-specific MAFs were extracted from 1000 Genomes Phase 3 VCFs.

### Analysis and visualization

Visualization of MAFs, and calculation and visualization of MAF differences between MVP and 1000 Genomes, was performed using R.

### HLA type predictions

4-digit HLA type predictions were generated for HLA-A and HLA-B from hard-called genotype data using HIBAG (Zheng et al. 2013). We chose the pre-fit Affymetrix Axiom UK Biobank Array 4-digit resolution model (https://hibag.s3.amazonaws.com/hlares_index.html), as the MVP genotyping array covers > 95% of this model’s training variants for both loci. We used the European model for individuals assigned to HARE group EUR and the multi-ethnic model for individuals assigned to HARE groups AFR, HIS, and ASN. Predictions were generated by calling the *predict()* function from HIBAG. Frequencies were calculated for each 4-digit allele by HARE group.

---

**Supplementary Figure 1.**
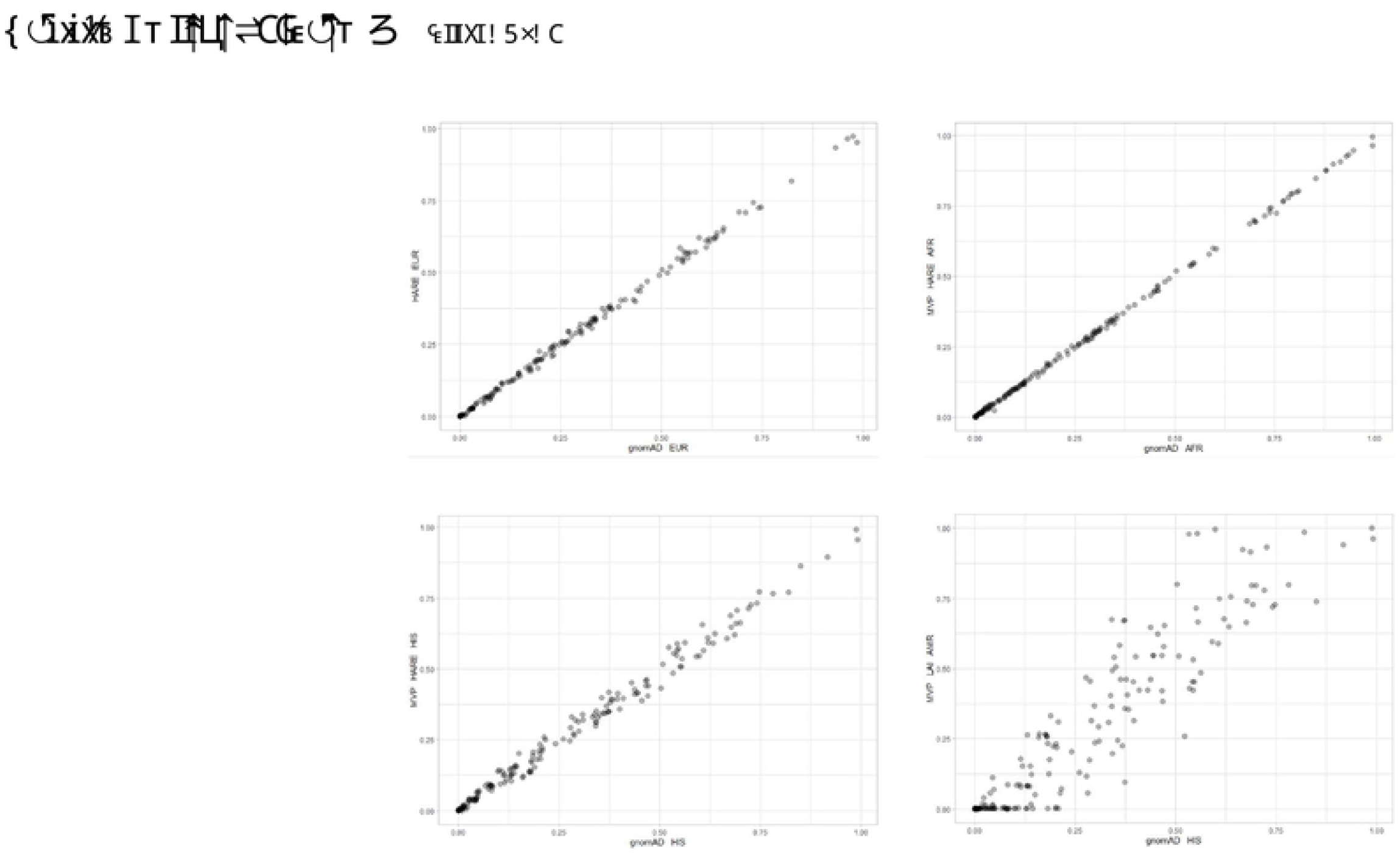
Allele frequency comparisons between gnomAD and MVP HARE groups for three groups (EUR, AFR, HIS). In the lower right we compare allele frequencies for gnomAD HIS and Local Ancestry Inference (LAI) derived allele frequencies for the AMR track of the HARE HIS group. We use three-way local ancestry deconvolution (EUR, AFR, AMR)

